# Depletion of voltage-dependent anion channel (VDAC) of *Toxoplasma gondii* affects multiple mitochondrial functions, but not calcium signalling

**DOI:** 10.1101/2020.10.07.330423

**Authors:** Natalia Mallo, Erica S. Martins Duarte, Stephan C. Baehr, Marco Biddau, Jana Ovciarikova, Mary-Louise Wilde, Alessandro D Uboldi, Leandro Lemgruber, Christopher J Tonkin, Jeremy G. Wideman, Clare R. Harding, Lilach Sheiner

**Author notes:** Corresponding –. Equal contribution (Clare R. Harding, Lilach Sheiner).

## Abstract

The Voltage Dependent Anion channel (VDAC) is a ubiquitous channel in the outer membrane of the mitochondrion with multiple roles in protein, metabolite and small molecule transport. In mammalian cells, VDAC, as part of a larger complex including the inositol triphosphate receptor, has been shown to have a role in mediating contact between the mitochondria and ER. We identify VDAC of the pathogenic apicomplexan *Toxoplasma gondii* and demonstrate its importance for parasite growth. We show that VDAC is involved in protein import and metabolite transfer to the mitochondria, but does not appear to modulate calcium (Ca2+) signalling. Further, depletion of VDAC resulted in significant morphological changes of the mitochondrion and ER, suggesting a role in mediating contacts between these organelles in *T. gondii*.

## Introduction

*Toxoplasma gondii* is the causative agent of toxoplasmosis and a member of the parasitic phylum Apicomplexa, which includes *Plasmodium*, the causative agent of malaria, and *Cryptosporidium*, which causes the diarrheal disease cryptosporidiosis. As a divergent eukaryote, *T. gondii* has a single, lasso-shaped mitochondrion which is required for respiration and metabolism 07/10/2020 14:36:00. To enable this, multiple small molecules, metabolites, and proteins must cross the two membranes of the mitochondrion. In many eukaryotes, from protists to mammals, transport across the outer mitochondrial membrane is mediated by a conserved, highly abundant porin named the Voltage-Dependent Anion Channel (VDAC) (Hodge and Colombini, 1997; Homblé et al., 2012; Wideman et al., 2013). VDAC mediates the passage of ions, nucleotides and metabolites across the outer membrane, and has been implicated in protein and tRNA import in to the mitochondria (Camara et al., 2017; Ellenrieder et al., 2019; Hodge and Colombini, 1997; Salinas et al., 2006). Beyond mediating transfer across the outer membrane, VDAC has also been identified as component of membrane contact sites between the mitochondria and endoplasmic reticulum (ER). VDAC clusters in domains of ER–mitochondria contacts (Rapizzi et al., 2002; Shoshan-Barmatz et al., 2004) where it interacts directly with the ER-resident inositol trisphosphate receptor (IP3R) via the chaperone Grp75 (Honrath et al., 2017; Szabadkai et al., 2006). This close apposition of membranes allows direct transfer of Ca^2+^ between the ER and the mitochondria, where the close association of the mitochondrial inner membrane Ca^2+^ uniporter (MCU) and associated protein MICU, on the inner membrane of the mitochondria allows uptake of Ca^2+^ into the organelle (De Stefani et al., 2016; Liao et al., 2015). This contact has been shown to have important roles in Ca^2+^ signaling in survival and proliferation in within mammalian cells (De Stefani et al., 2016; Rieusset et al., 2016; Rizzuto et al., 2009)

In *T. gondii*, the ER is considered the major Ca^2+^ store and key aspects of the parasite’s life cycle, such as host-cell invasion and gliding motility, are initiated and regulated by Ca^2+^ release, largely thought to be from the ER (Borges-Pereira et al., 2015; Lourido and Moreno, 2015; Lovett and Sibley, 2003; Wetzel et al., 2004). To support this, parasite motility can be blocked by inhibition of the second messengers inositol 1,4,5-trisphosphate (IP_3_), and cyclic ADP-ribose (cADPR), which act on currently undefined, ER-resident receptors (Lovett et al., 2002; Triana et al., 2018). However, there is evidence that other organelles, including the mitochondrion and acidocalcisomes, also play a role in Ca^2+^ storage in Apicomplexa (Miranda et al., 2010; Nagamune et al., 2007; Rohloff et al., 2011), and the relative importance of these potential stores in modulating invasion and motility remains unknown.

Although the function and composition of the *T. gondii* mitochondrion has been a growing focus of study in recent years, little is known about how molecules are transported across the mitochondrial membranes of apicomplexan parasites. Here, we identified and characterized VDAC of *T. gondii*. We found that VDAC is important for fitness, and its depletion results in changes in mitochondrial and ER morphology in concert with reduced ER-mitochondrion contacts, suggesting for the first time the existence of mitochondrial-ER contact sites in this parasite. Interestingly, this was not accompanied by changes in cytosolic Ca^2+^ or in processes that depend on Ca^2+^ signalling, pointing to functional divergence between parasites and mammalian cells.

## Results

### *T. gondii* VDAC is required for parasite growth

The gene encoding *T. gondii* VDAC (TGME49_263300, XP_002365430.1) was previously identified (Wideman et al., 2013). Using HHPRED, TGME49_263300 had high (E-value = 2.2 e^-39^) structural homology to mammalian VDAC, including a conserved glutamate at position 16 which has previously been shown to have a role in voltage conductance (**Fig. S1A)** (Shuvo et al., 2016). Gene ontology suggested anion transport and voltage-gated channel activity, and data from the recent *T. gondii* protein atlas predicted mitochondrial localization, together supporting VDAC orthology (Barylyuk et al., 2020; Gajria et al., 2008). We thus named TGME49_263300, VDAC. VDAC is conserved between Apicomplexa and its phylogenetic analysis sequences recapitulated the phylogenetic distribution of the corresponding organisms (**Fig. S1B**). To localise VDAC within the parasite, the predicted mRNA sequence was cloned into a *T. gondii* expression vector with a strong promotor and N-terminal Myc epitope tag (Harding et al., 2016) and parasites transiently expressing this exogenous copy were imaged. Immunofluorescence analysis demonstrated co-localisation of Myc-VDAC with the mitochondrial outer membrane marker TgMys (Ovciarikova et al., 2017) (**Fig. 1A**). Interestingly, Myc-VDAC was not distributed smoothly along the mitochondria but instead was concentrated in specific patches. This non-uniform localization has been observed in mammalian cells where VDAC form clusters at ER-mitochondrial contact sites (Rapizzi et al., 2002).

**Figure 1.**
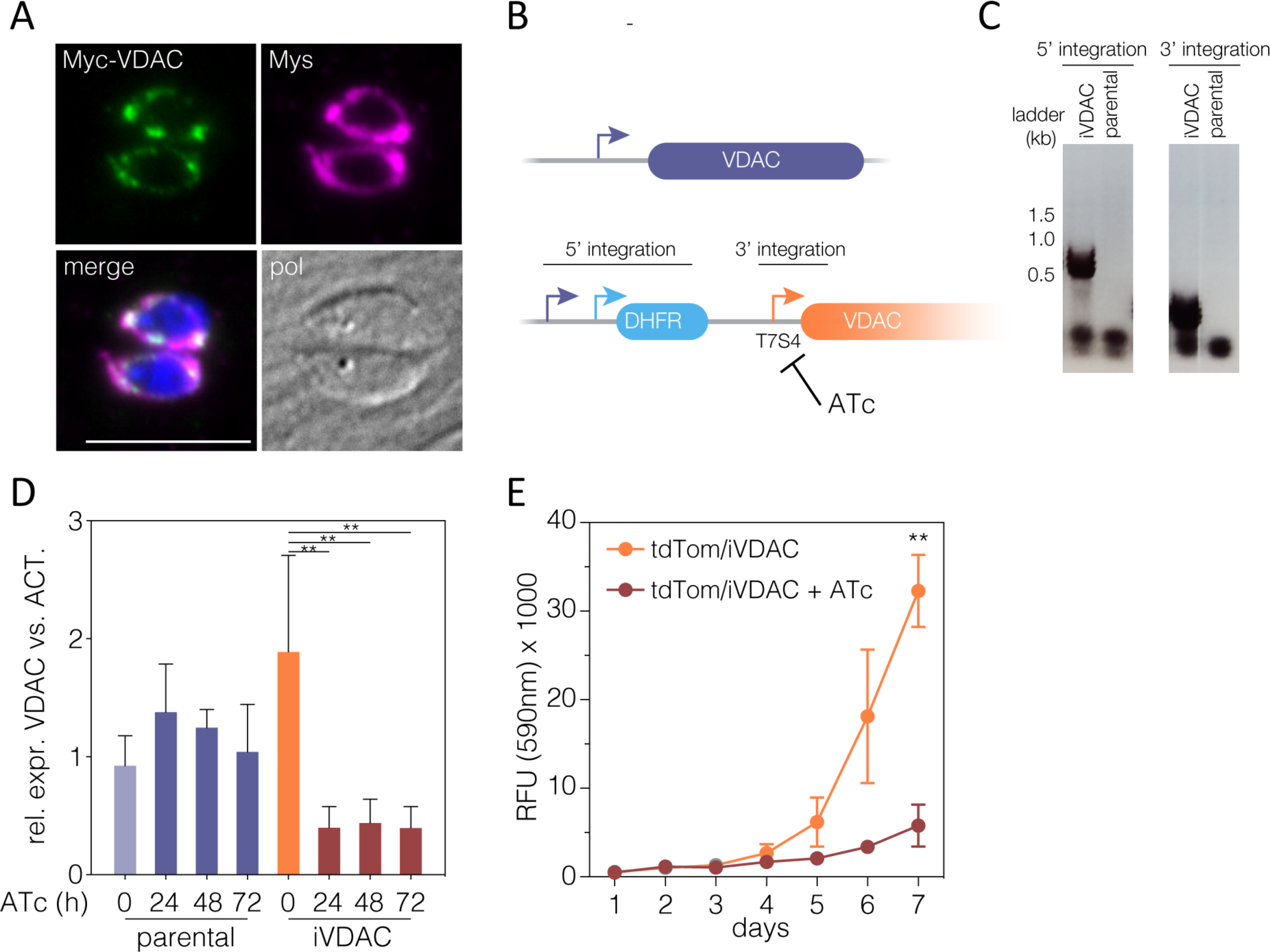
Downregulation of VDAC leads to decrease in parasite growth. **A)** Overexpression of Myc-VDAC (green) co-localized with anti-Mys, a mitochondrial membrane marker. Scale bar 5 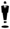m. **B)** Schematic representation of the promoter replacement strategy used to create iVDAC parasite line. **C)** PCR to confirm integration of the regulatable promoter and the DHFR selectable marker cassette, expected PCR products depicted in (b). **D)** Expression of *VDAC* relative to *actin* measured by RT-qPCR over ATc treatment. Bar represents mean 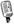SD ** p < 0.005, statistical analysis performed with one-way ANOVA with Turkey correction, *n* = 3 independent experiments. **E)** Fluorescence of iVDAC parasites stably expressing tdTomato 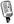ATc over time, results are mean fluorescence (arbitrary units) 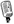SD. ** *p* = 0.005, Student’s t test corrected for multiple tests using Holm-Sidak, n = 3.

Results from genome-wide CRISPR screens suggested that VDAC was important for fitness in *T. gondii* (Sidik et al., 2016). To examine the role of VDAC in the lifecycle in more detail, we constructed a conditional knockdown line by replacing the VDAC promoter with a T7S4 promoter which is repressed upon addition of anhydrotetracycline (ATc) (Lacombe et al., 2019; Sheiner et al., 2011) (**Fig. 1B**). Integration of the regulatable promotor was confirmed by PCR (**Fig. 1C**). Downregulation of the corresponding mRNA upon ATc addition was assessed by RT-qPCR, demonstrating an 80% reduction in mRNA levels at 24 h post treatment (**Fig. 1D**). Growth of fluorescent iVDAC parasites was significantly inhibited by seven days post ATc addition (**Fig. 1E**), demonstrating that VDAC is important for parasite fitness.

### Depletion of VDAC leads to disruption of mitochondrial morphology

It has been shown that disruption of VDAC can cause changes in mitochondrial morphology (Ferecatu et al., 2018; Park et al., 2010). Previously, we defined three main mitochondrial morphologies detected by immunofluorescence microscopy in wild type intracellular *Toxoplasma* tachyzoites (Ovciarikova et al., 2017). VDAC depletion resulted in three additional morphologies, which we named connected (for mitochondria that are connected between a number of parasites), broken and ball-shaped, and which we scored from microscopy images (**Fig. 2A**). Addition of ATc to the parental line did not affect mitochondrial morphology, which presented as mostly lasso or open lasso shapes as previously reported (Lacombe et al., 2019; Ovciarikova et al., 2017) (**Fig. 2B**). However, treatment of the iVDAC parasite line revealed a significant increase in abnormal morphologies at both 48 and 72 h post ATc addition (**Fig. 2B**).

**Figure 2:**
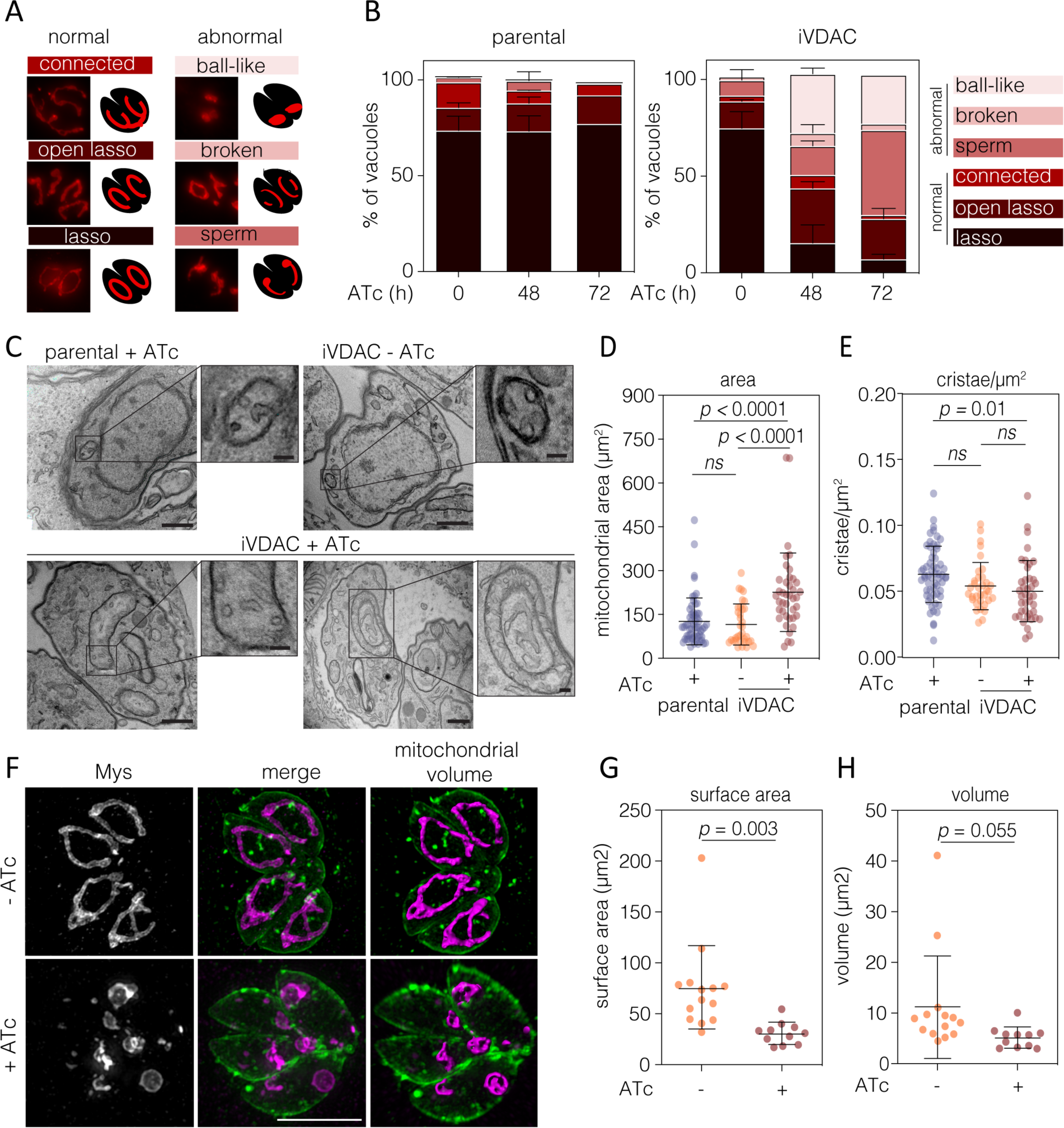
Depletion of VDAC results in mitochondrial morphological defects. **A)** Mitochondrial morphology of intracellular parasites was scored as indicated. **B)** Quantification of mitochondrial morphology from parental and iVDAC lines at indicated timepoints of ATc treatment. 100 vacuoles from two independent experiments were quantified, results are mean + SD. Between ‘normal’ and ‘abnormal’ morphologies, one-way ANOVA with Dunnettes correction were performed across time points. No significant changes were seen in the parental lines upon ATc treatment, but there were significantly more abnormal morphologies in iVDAC at 24 (*p* = 0.012) and 48 h (*p* = 0.039) post ATc addition. **C)** Representative TEM images of parental parasites and iVDAC parasites, untreated or treated with ATc for 48 h, scale bars 500 nm. Insets showing detail of mitochondrion structures, scale bars 100 nm. Quantification of mitochondrial area **(D)** and number of cristae/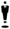m^2^ **(E)** from EM images. At least 30 parasites quantified per condition from n = 2 independent experiments, results are mean 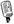SD, *p* values from one-way ANOVA with Tukey correction. **F)** SR-SIM projections of mitochondrion (Mys, magenta) and plasma membrane (SAG1, green) of iVDAC parasites, untreated and treated with ATc for 48 h. Volume projections of mitochondria shown, where substantial changes in mitochondrial morphology can be seen. Scale bar 5 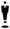m. Quantification of mitochondrial surface area **(G)** and volume **(H)** calculated from volume projections, of at least 10 vacuoles containing 2 parasites. Results mean 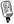SD, *p* values from Student’s t-test.

To investigate the morphology of the mitochondrion in more detail, transmission electron microscopy was used. In agreement with the above observation, there were no apparent effects of ATc on the parental mitochondria. In contrast, at 48 h post ATc treatment, the iVDAC mitochondrion appeared larger and occasionally contained vesicular structures (**Fig. 2C**). Quantification of mitochondrial area from TEM images demonstrated that in the sections imaged, the regions of mitochondria had significantly greater area in the iVDAC line after ATc treatment (**Fig. 2D**). VDAC depletion has been shown to affect cristae morphology in mouse muscle fibre (Baghel and Thakur, 2019) and previously, depletion of a mitochondrial protein in *T. gondii* has resulted in a change of the density of mitochondrial cristae (Huet et al., 2018), however, depletion of VDAC in *T. gondii* did not change the density of cristae within the mitochondria (**Fig. 2E**).

TEM sections were selected based on the presence of mitochondria and so do not represent the total mitochondrial volume of the parasites. To determine how the total volume of mitochondrion changed upon depletion of VDAC, we quantified the surface area and total volume of mitochondria per parasite, using 3D super resolution-structured illumination microscopy (SR-SIM). Z-stacks of vacuoles containing 2-4 iVDAC parasites at 48 h post ATc addition stained with anti-Mys (mitochondrion) and anti-SAG1 (parasite plasma membrane) were collected (**Fig. 2F**). The volume and surface area of mitochondria were automatically quantified from 3D rendered projections of the anti-Mys signal. Depletion of VDAC resulted in a 59 % decrease in mitochondrion surface area (**Fig. 2G**) and a 55 % decrease in mitochondrion volume at 48 h (**Fig. 2H**). These results demonstrate that loss of VDAC results in a significant alteration in gross mitochondrial morphology.

### VDAC depletion leads to alterations in mitochondrial metabolism and protein import

Due to the growth defect and the significant alterations in mitochondrial morphology, we wanted to determine whether mitochondrial physiology was altered upon VDAC depletion. Parasite mitochondrial membrane potential (ΔΨm) was assessed using flow cytometry of parasites stained with the fluorescent probe JC-1 (**Fig. 3A**) as previously described (Brooks et al., 2010). As expected, treatment with valinomycin (val) resulted in a decrease in the ratio of red to green fluorescence, indicating a loss of membrane potential (Brooks et al., 2010). In contrast, depletion of VDAC for 48 h did not result in a significant change in mitochondrial membrane potential (**Fig. 3A**).

**Figure 3:**
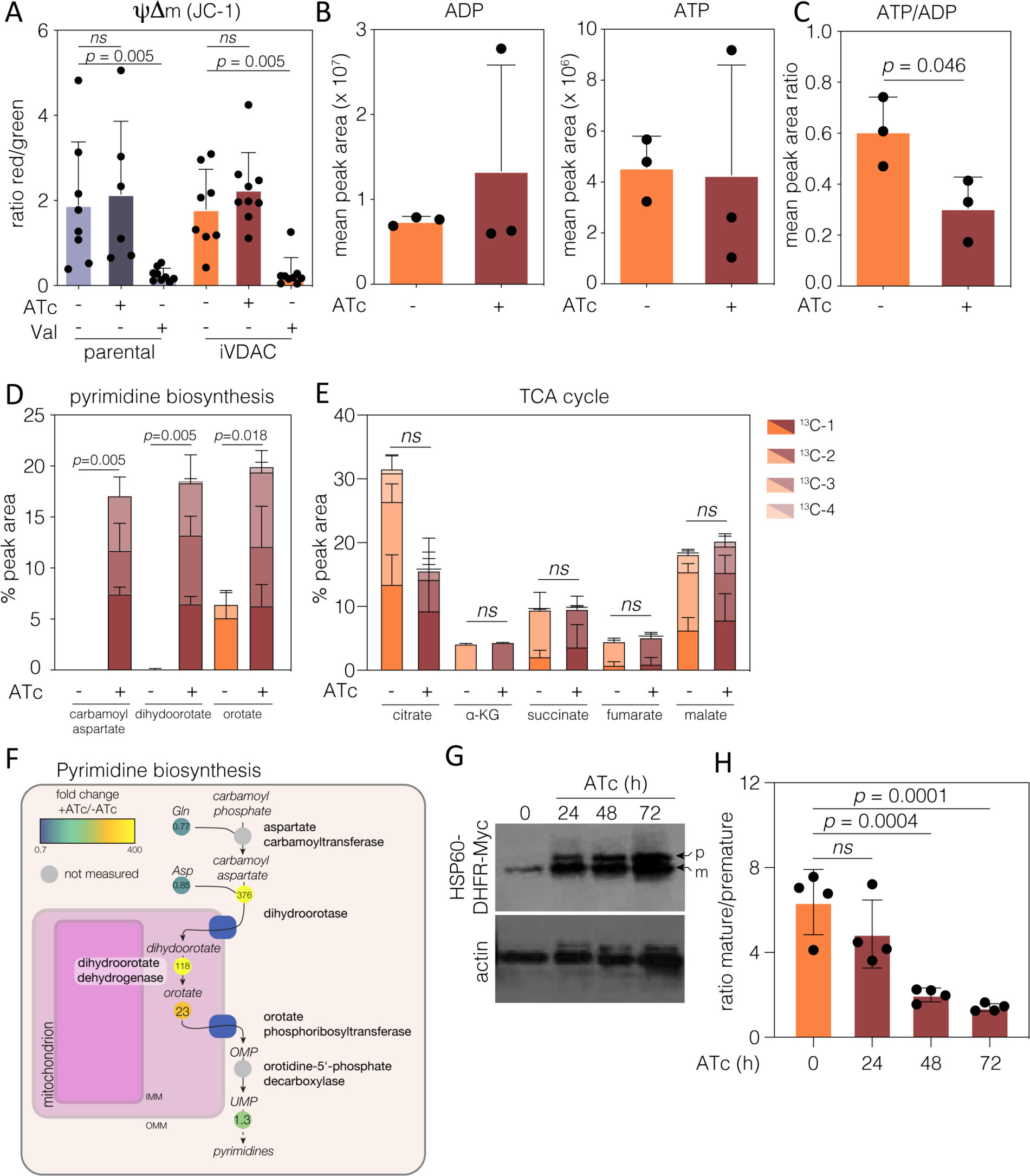
Mitochondrial functions are mildly affected by depletion of VDAC. **A)** Mitochondrial membrane potential quantified using JC-1 probe. Results show mean ratio of red/green fluorescence events 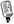SD, as recorded by flow cytometry. Valinomycin (Val) was included as a membrane potential depolarizing control. *p* values from Student’s t-test (n = at least 5 independent experiments). **B)** Quantification of ADP and ATP levels from labelled metabolomics, values are combination of labelled and unlabeled peak areas. Points are mean of the peak recorded area 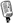SD, n = 3. **C)** Ratio between ADP and ATP levels, *p* values from Student’s t test, n *=* 3. Quantification of selected labelled metabolites in the pyrimidine biosynthesis pathway **(D)** and the TCA cycle **(E)** from iVDAC parasites untreated and at 48 h post ATc addition. Carbamoyl aspartate and dihydroorotate were almost undetectable in untreated parasites. Bar represents mean + SD of each species of labelled metabolite, n = 3. **F)** *T. gondii* pyrimidine pathway showing fold change in metabolite levels upon treatment with ATc, along with the predicted localization of key enzymes in the pathway. Dihydroorotate dehydrogenase (DHODH) is found on the outer face of the inner mitochondrial membrane. **G)** Protein import assays were performed after transient transfection of the iVDAC line with HSP60L-DHFR-myc and treatment with ATc for the indicated time. Premature (p) and mature (m) bands can be identified by western blot, actin used as loading control. Accumulation of premature form can be seen as early as 24 h post ATc treatment. Representative of n = 4 experiments. **H)** Densitometry obtained from bands shown in (**G**) regarding mature/premature ratio after ATc treatments, results mean 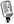SD, *p* values from one-way ANOVA with Dunnets correction, n = 4.

In mammalian cells, VDAC is the major ATP and ADP transporter across the mitochondrial outer membrane, and depletion of VDAC isoforms results in a decrease in the cellular ATP/ADP ratio (Krammer et al., 2015; Maldonado et al., 2013). Untreated and 48 h ATc-treated iVDAC parasites were labelled with isotope-labelled ^13^C-U-D-glucose and total ATP and ADP levels were quantified by LC-MS (**Fig. 3B**). No significant changes in ADP or ATP levels were seen upon VDAC depletion, however we did observe a small (*p* = 0.046, unpaired *t* test) decrease in the ATP/ADP ratio (**Fig. 3C**), suggesting that VDAC may have a role in nucleotide transport in *T. gondii*, although it is possible that the residual VDAC expression present allows for some level of nucleotide transport.

In addition to ATP/ADP, VDAC is responsible for transportation of numerous small metabolites across the mitochondrial outer membrane (Hodge and Colombini, 1997; Lee et al., 2011; Pusnik et al., 2009). To determine how VDAC depletion affected global parasite metabolism, we analysed the ^13^C-U-D-glucose labelled metabolites further. We observed a dramatic accumulation of carbomyl aspartate, dihydroorotate and orotate (**Fig. 3D**) after VDAC depletion. These metabolites are intermediates in the pyrimidine biosynthesis pathway (**Fig. 3F**). Interestingly, both carbomyl aspartate and dihydroorotate were undetectable in the untreated iVDAC line, while orotate was present at a low level. We did not see changes in levels of pyrimidines upon VDAC depletion (results not shown), however *T. gondii* is known to have a pyrimidine salvage pathway (Donald and Roos, 1995), which might explain the steady pyrimidine level. These results suggest that depletion of VDAC leads to dysregulation of the pyrimidine biosynthesis pathway. To confirm that addition of ATc did not cause a global change in mitochondrial metabolism, we also examined levels of the TCA cycle intermediates, citrate, ✓-ketoglutarate, succinate, fumarate and malate, which are expected to have specialised transports. We saw no significant changes in levels upon ATc treatment (**Fig. 3E**), suggesting that addition of ATc did not induce global changes to mitochondrial metabolism.

Beyond small molecules, VDAC also has a role in mitochondrial protein import in both plants and yeast through interacting with components of the translocon machinery (Ellenrieder et al., 2019; Salinas et al., 2006). To determine if depletion of VDAC affected mitochondrial protein import in *T. gondii*, we assessed the maturation of the mitochondrial-targeted marker HSP60L-DHFR-Myc (van Dooren et al., 2016) through its transient expression in iVDAC parasites (**Fig. 3G**). A similar technique has been used successfully to assess protein import into the apicoplast and mitochondria in *T. gondii* (Sheiner et al., 2015, 2011; van Dooren et al., 2016). Depletion of VDAC caused a significant accumulation of the premature form of the protein by 48 h post ATc addition (**Fig. 3H**), suggesting that protein import into the mitochondrion is disrupted by depletion of VDAC.

### Putative mitochondrion-ER contact sites are reduced upon VDAC depletion

In mammalian cells, VDAC, along with ER and cytosolic components, are involved in Ca^2+^ regulation at sites of mitochondrial-ER juxtaposition, suggested to be membrane contact sites (MCS) (Szabadkai et al., 2006). To determine if similar sites of membrane apposition might be present between the mitochondrion and the ER in *T. gondii*, we analysed random thin sections of the parental parasite line. Morphologically, we defined membrane apposition as a constant distance of less than 30 nm between the two organelles over stretches of at least 100 nm (Ovciarikova et al., 2017). Using these parameters, putative contacts between the mitochondria and IMC were seen in 56.4 % of mitochondrial sections (taken from 94 parasites), as previously reported (Jacobs et al., 2020; Ovciarikova et al., 2017). Membrane apposition between the mitochondria and ER were seen in 64.4 % of mitochondria analysed (**Fig. 4A**), implying that close contact between these organelles is frequent in *T. gondii*. Sections of iVDAC parasites, treated and untreated with ATc, revealed that depletion of VDAC led to a reduction of both IMC/mitochondria (from 40.2% to 30.0% of mitochondria) and ER/mitochondria contacts (from 64.8% of mitochondria to 44.5%) (**Fig. 4B**). Thus, VDAC depletion results in reduction of mitochondrial contacts with both the IMC and the ER, with a higher impact on ER-mitochondrial membrane apposition.

**Figure 4.**
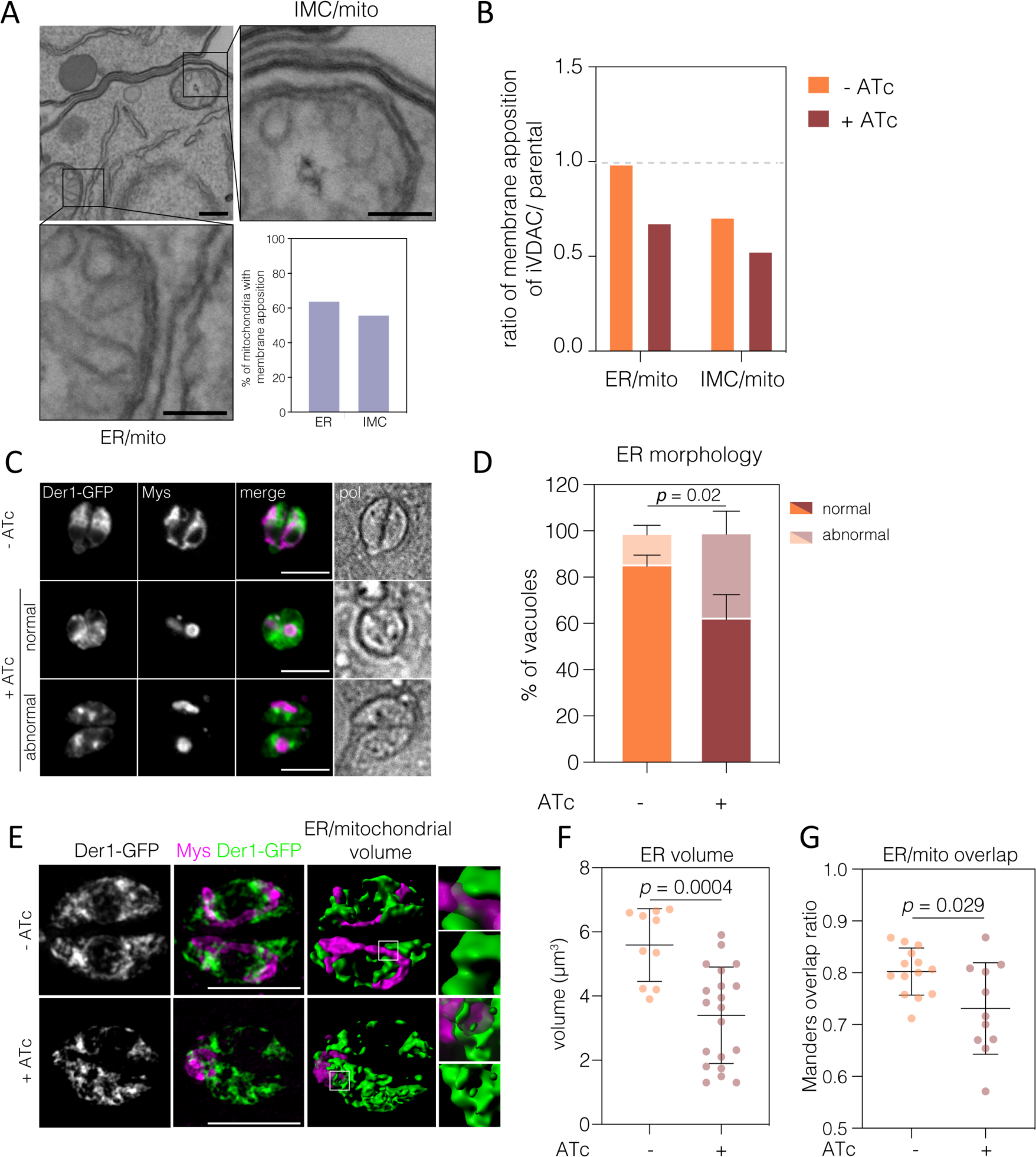
Depletion of VDAC leads to changes in ER morphology. **A)** Representative TEM of putative ER/mitochondria and proposed IMC/mitochondria membrane contact site (MCS) from parental parasites. Scale bars 500 nm, insets 100 nm. Graph shows the proportion of parental-line parasites with visible areas of ER and IMC membrane apposition sites, results from 94 sections. **B)** The proportion of areas of membrane apposition from iVDAC parasites with or without ATc treatment, normalised to parental parasites. At least 88 sections quantified per condition. **C)** iVDAC parasites transiently expressing the ER marker Der1-GFP were ATc treated for 48 h, fixed and ER morphology examined. Upon VDAC depletion, the ER of many parasites lost its normal morphology and rounded up into small foci within the parasite. Scale bars 5 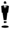m. **D)** Quantification of ER morphology, scored as normal or abnormal. Results mean + SD, n = 3. Significant increase in abnormal morphology after treatment, *p* = 0.019, Student’s t test. **E)** Representative 3D reconstructions of ER (Der1-GFP, green) and mitochondria (Mys, magenta) in iVDAC parasites at 48 h post ATc addition. Scale bars 5 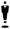m. **F)** The volume of parasite ER was calculated from untreated and treated parasites. Points represent individual parasite ER with lines at mean, 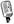SD. Significance from Student’s t-test. **G)** The ratio of overlap of the ER (Der1-GFP) with the mitochondria (anti-Mys) was quantified from SR-SIM images. Box plots are mean with whiskers at 10 – 90 %. Significance from Welches t-test, ratios calculated from at least 11 vacuoles.

To examine how the endoplasmic reticulum was affected by VDAC depletion, iVDAC parasites were transiently transfected with a construct expressing the ER-resident Derlin1 (Der1) fused to GFP (Agrawal et al., 2009). We found that depletion of VDAC for 48 h resulted in significant changes in the localization and distribution of the ER (**Fig. 4C**). In the absence of VDAC, the ER morphology changed from a perinuclear, reticular network found throughout most of the cytoplasm in untreated parasites, to the accumulation of Der1-GFP in foci and loss of the spread, perinuclear distribution. 48 % of VDAC-depleted parasites displayed this abnormal ER localisation, as opposed to only 13% of the untreated parasites (**Fig. 4D**).

On the basis of the profound effect of VDAC depletion on mitochondrial and ER morphology, we examined other parasitic organelles. However, we saw no changes in the parasite pellicles (known as the inner membrane complex, (IMC)) (**Fig. S2A**), the secretory micronemes (**Fig. S2B**) and rhoptries (**Fig. S2C**), or the parasite’s plastid (apicoplast) (**Fig. S2D**) at 48 h post ATc treatment.

To investigate the effect of VDAC depletion on the ER in more detail, we acquired SR-SIM images of iVDAC parasites expressing Der1-GFP (**Fig. 4E**). Upon depletion of VDAC, we observed a significant decrease in the volume of parasite ER signal in reconstructed 3D projections (**Fig. 4F**). We also observed a decrease in the proportion of ER/mitochondrial fluorescence signal overlap (**Fig. 4G**). These results suggest that the defects of the mitochondrial morphology coincide with defect in both the distribution of the ER, and the frequency of contacts between the mitochondrion and ER.

Together, these results suggest that MCS may be present between the mitochondrion and ER, and that depletion of VDAC leads to both a decrease in MCS and a change in the morphology of the ER, but not other parasite organelles.

### VDAC depletion does not change cytosolic Ca^2+^ in *T. gondii*

There is biochemical evidence that the ER is a major Ca^2+^ store in *T. gondii* (Nagamune et al., 2007), however the role of the mitochondrion in Ca^2+^ storage and release in these parasites is still unknown. We first analysed cytosolic [Ca^2+^] levels using genetically encoded Ca^2+^ sensor GCaMP6 normalized to mCherry fluorescence as previously described (Stewart et al., 2017). Using this method, we saw no change in GCaMP6 fluorescence upon depletion of VDAC (**Fig. 5A**). To investigate the response to Ca^2+^ store release in the absence of VDAC, GCaMP6 expressing parasites were stimulated with increasing concentrations of either the Ca^2+^ ionophore ionomycin (IO) or the specific phosphodiesterase inhibitor 5-benzyl-3-isopropyl-1H-pyrazolo[4,3-d]pyrimidin-7(6H)-one (BIPPO), which increases levels of cGMP leading to activation of PKG-dependent signalling and a downstream rise in cytosolic [Ca^2+^] (Stewart et al., 2017). In both cases, treatment with the drug resulted in a concentration-dependent increase in GCaMP6 signal, demonstrating effective release of Ca^2+^ stores (**Fig. 5B**). However, no difference in GCaMP6 signal was seen upon depletion of VDAC, indicating that VDAC likely does not have a role in stimulated Ca^2+^ release under these conditions. However, as the ER and acidocalcisome are the major Ca^2+^ stores within the cell (Contreras et al., 2010; Rohloff et al., 2011; Triana et al., 2018), changes in Ca^2+^ handling from the mitochondria may be too small, localized or transient to be detected by changes in total cytosolic GCaMP6 fluorescence.

**Figure 5:**
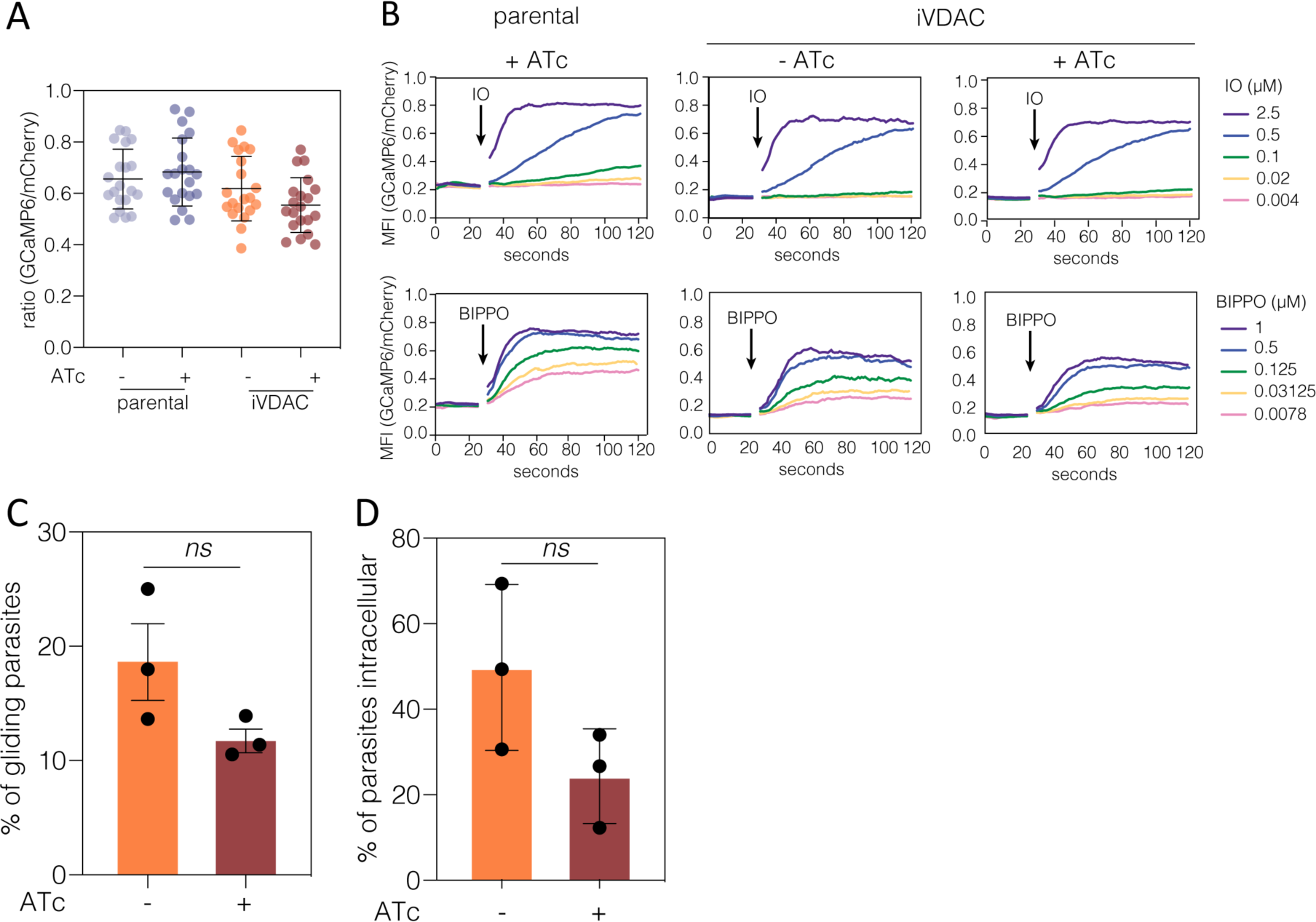
VDAC depletion affects Ca^2+^ induced gliding motility but has no effect on cytosolic Ca^2+^. **A)** Parental and iVDAC parasites expressing mCherry and GCaMP were analysed by flow cytometry and the GCaMP/mCherry ratio calculated in resting parasites. No significant difference could be seen in the ration upon depletion of VDAC. Bars represent mean 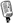SD from 4 independent experiments. **B)** GCaMP/mCherry ratio was calculated after stimulation with various concentrations of ionomycin or BIPPO for 120 s. No differences could be seen upon stimulation of Ca^2+^ release upon VDAC depletion. **C)** The percentage of iVDAC parasites with trails was quantified. Bars represent mean 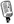SD, ns – no significant difference, Students t test, n = 3. **D)** Percentage of iVDAC parasites internalized after 20 minutes, ns – no significant difference, Students t test, n = 3.

Both gliding motility and invasion are regulated by transient changes in cytosolic Ca^2+^ levels. To determine the effect of VDAC depletion on these processes, we quantified the percentage of gliding (**Fig. 5C**) and invaded (**Fig. 5D**) parasites, after 72 h ATc treatment. We saw no significant change in either gliding or invasion, confirming that depletion of VDAC is unlikely to alter intracellular Ca^2+^ mobilization.

In light of previous work associating VDAC with Ca^2+^ mobilization at ER-mitochondrion contact sites (Honrath et al., 2017; Min et al., 2012; Rieusset et al., 2016; Szabadkai et al., 2006), our results showing no change in Ca^2+^ mobilisation were unexpected. In animals, VDAC mediates Ca^2+^ transfer from the ER to the mitochondria through association with the ER-localised IP3R via Grp75 in the cytosol (Min et al., 2012; Szabadkai et al., 2006). The mitochondrial Ca^2+^ uniporter (MCU) and associated protein MICU interact with VDAC and are also required for Ca^2+^ transfer across the mitochondrial membranes (De Stefani et al., 2016; Rizzuto et al., 2009). While VDAC and Grp75 are ubiquitous across eukaryotes (Wideman et al., 2013), MCU, MICU, and IP3Rs (or the paralogous ryanodine receptors (RYR), previously been reported to have been lost from multiple lineages, including some apicomplexans (Bick et al., 2012). On the basis of a number of new genomes becoming available, we re-investigated the presence of MCU/MICU and IP3R/RYR3 in alveolates (**Fig. 6**) and wider eukaryotes (**Fig. S3**). We find no evidence for MCU, MICU or IP3R/RYR3 in *T. gondii, P. falciparum*, or *Cryptosporidium* sp. (**Fig. 6**). However, we identified orthologues of MCU and MICU in *Besnoitia besnoiti* and *Cystoisospora suis*, parasites closely related to *T. gondii*. In addition, we find components in chromerid algae closely related to apicomplexans, e.g. MCU, MICU, and IP3R/RYR are present in *Vitrella brassicaforma* and MCU and MICU are found in *Chromera velia* **(Fig. 6)**. These data demonstrate that genes involved in mitochondrial Ca^2+^ transfer in mammals have been independently lost at least 19 times through evolution, including in apicomplexans.

**Figure 6:**
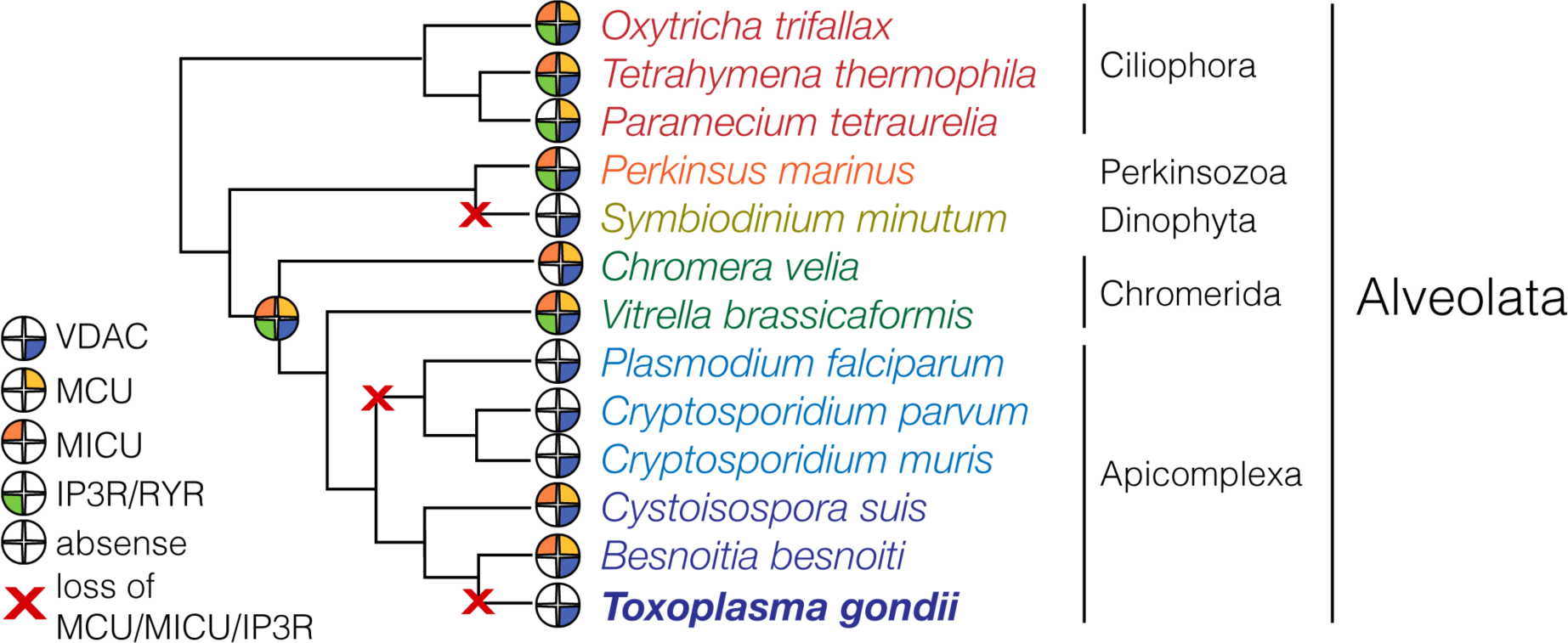
VDAC is ubiquitous across alveolates, but components of the calcium contact, MCU, MICU, and IP3R are absent from most apicomplexans. Putative orthologues of VDAC, MCU, MICU, and IP3R were identified by the reciprocal BLAST method. Filled pies indicate presence of a particular orthologue. Historical losses of all of MCU, MICU, and IP3R are indicated with red Xs. Although VDAC was not identified in *Symbiodinium minutum* (likely due to the incomplete database), it was identified in a closely related species of *Symbiodinium* -*S. microadriaticum*.

## Discussion

Here we describe a *T. gondii* homolog for the ubiquitous mitochondrial porin, VDAC. As the most abundant pore in the outer mitochondrial membrane across phyla, VDAC performs an important role in allowing passage of macromolecules and metabolites between the cytosol and intermembrane space. In *T. gondii*, depletion of VDAC resulted in subtle changes in metabolite abundance and has no significant effect on the mitochondrial membrane potential or Ca^2+^ signalling. Instead, dramatic changes in mitochondrial and ER morphology were observed, in parallel with reduced protein import and impaired parasite growth. These results demonstrate the conserved roles of VDAC across eukaryotes, while highlighting some important changes in the Apicomplexa.

One of the canonical roles of VDAC is as pore for nucleotide transport across the outer mitochondrial membrane. In several systems, depletion of VDAC leads to a decrease in the ATP/ADP ratio (Maldonado et al., 2013; Maldonado and Lemasters, 2014) due to inhibition of the flow of ATP and ADP across the mitochondrial membrane. In *T. gondii* we see a similar phenotype; while absolute levels of cellular ATP and ADP do not differ substantially, the ratio of ATP/ADP decreases, which suggests a role for VDAC in mediating nucleotide transport across the mitochondrion in these parasites. Given that ∼20 % VDAC expression is maintained under ATc, it is possible that the residual VDAC is able to perform some of this function under these conditions, thus explaining the relatively mild defect at the time point tested. VDAC also acts as a pore for other metabolites including amino acids and other metabolic intermediates (Ellenrieder et al., 2019; Hodge and Colombini, 1997; Pusnik et al., 2009). Interestingly, we found that depletion of VDAC leads to a large accumulation of intermediates in the pyrimidine biosynthesis pathway. Pyrimidine synthesis occurs in the cytosol, with the exception of the conversion of dihydroorotate to orotate, catalysed by dihydroorotate dehydrogenase (DHODH), which is localized to the inner mitochondrial membrane in *T. gondii* (Triana et al., 2012). In *Plasmodium*, chemical inhibition of DHODH leads to accumulation of carbamoyl aspartate and dihydroorotate (Creek et al., 2016). We suggest that dihydroorotate, and possibly orotate, requires transport via VDAC across the outer mitochondrial membrane and in the absence of VDAC, intermediates in the pathway accumulate. Depletion of VDAC does not appear to affect the function of the TCA cycle which relies on specialized solute transporters (van Dooren et al., 2006), demonstrating that the effect of VDAC depletion is specific to the pyrimidine pathway. The pyrimidine pathway is of major interest in *Plasmodium* where DHODH has emerged as an important drug target (Hoelz et al., 2018; Painter et al., 2007) and this potential role of VDAC will merit further study.

VDAC has been shown to be linked to protein import across the outer mitochondrial membrane in yeast and plants (Ellenrieder et al., 2019; Salinas et al., 2006). In *T. gondii*, depletion of VDAC led to a defect in the import of a mitochondrial localised HSP60L, previously used to demonstrate protein import defect upon depletion of components of the TOM complex (van Dooren et al., 2016). Interestingly, it was previously suggested that, although the TOM complex is essential for the parasite, some protein import could occur in its absence (van Dooren et al., 2016). It is possible that VDAC is involved in this bypass. However, VDAC depletion also leads to morphological collapse of the mitochondrion, and this change in morphology may indirectly inhibit protein import.

In plant cells VDAC is also thought to be required for import of tRNAs (Salinas et al., 2014, 2006). The mitochondrial genome of *T. gondii* is severely reduced and encodes only subunits of the electron transport chain and rRNAs, which require tRNA import for translation (Esseiva et al., 2004; Lacombe et al., 2019; Namasivayam et al., 2020; Pino et al., 2010). However, we do not see any changes in the membrane potential of the mitochondria upon VDAC depletion, providing indirect evidence that the electron transport chain remains functional, thus likely no defect in tRNA import occurs. This is in agreement with results from *Trypanosoma* which does not require VDAC for tRNA import (Pusnik et al., 2009). The mechanism for tRNA import into the mitochondrion of *T. gondii* remains unknown and highlights the diverse strategies evolved to fulfil this essential function.

In *T. gondii*, close apposition of organelles has been noted, including between the mitochondrion and IMC (Jacobs et al., 2020; Ovciarikova et al., 2017), and the mitochondrion and apicoplast (Kobayashi et al., 2007; Nishi et al., 2008), however the molecular identity of many of the players is only beginning to be understood (Jacobs et al., 2020). Based on TEM images of wild type parasites, the mitochondrion of *T. gondii* appears to form frequent, close associations with the ER. VDAC is a natural candidate to mediate this interaction, based on results from other organisms, and in *T. gondii* depletion of VDAC leads to significant and specific changes in both mitochondrion and ER morphology. This is in contrast to LMF1, a mitochondrial protein whose deletion leads to a change of mitochondrial morphology, but no change to ER distribution (Jacobs et al., 2020). These results suggest that VDAC is involved in interactions between the mitochondrion and the ER, which are important to maintain the distribution of both organelles throughout the parasite. In mammalian cells the distribution of organelles is also mediated and supported by the tubulin, and to a lesser extent, actin cytoskeletons (Barlan and Gelfand, 2010; Frederick and Shaw, 2007). However, in *T. gondii* the tubulin cytoskeleton is involved in maintaining cell shape but does not pass through the body of the cell (Morrissette and Sibley, 2002), and alterations in parasite actin do not appear to affect the mitochondrion or ER morphology (Tosetti et al., 2019). This may suggest that MCS in *T. gondii* have a more important role in maintaining organelle distribution than in other species, as suggested upon the mitochondrial-IMC contacts description (Ovciarikova et al., 2017).

The mammalian contact involving VDAC, GRP75 and IP3R interacts with MCU and MICU to regulate Ca^2+^ mobilisation. Our phylogenetic profiling confirmed MCU, MICU, and IP3R as ancestral eukaryotic genes. Each major lineage evolved from an ancestor with a full complement of these proteins; however, we report no fewer than 19 independent losses of all three components (Figure S3). The co-occurrence of these proteins in many lineages is suggestive of their functional overlap, potentially as mediators of Ca^2+^ signalling as proposed for mammals. Although these proteins are ancestral and critical components of eukaryotes, our data suggest that either A) compensatory mechanisms exist that can take the role of this system when lost, B) the system can be readily replaced by a novel mechanism through functional innovation or horizontal gene transfer, or C) that the well-studied functions of these proteins in animals represent a lineage-specific adaptation. From our data on VDAC we cannot distinguish these options; however future work on determining the composition of mitochondrial/ER contacts in Apicomplexa may help to answer these questions.

In summary, our results give the first indication that the broadly conserved outer mitochondrial porin VDAC is present and functional in *T. gondii*. VDAC appears to have a role in protein and metabolite transfer and is required to maintain the morphology of both the mitochondria and the ER. The study of membrane contact sites in the context of this divergent parasite is a new area with likely important implications in signalling regulation during the lytic lifecycle. Our observed multiple losses of components of the mammalian Ca^2+^ signalling contact along with the finding that VDAC depletion does not lead to defect in Ca^2+^ signalling, highlight divergence between parasite and host in this critical pathway.

## Acknowledgments

This work is supported by BBSRC BB/N003675/1 grant and Wellcome Investigator Award 217173/Z/19/Z to L.S. L.S is a Royal Society of Edinburgh Personal Research Fellow. C.R.H holds a Sir Henry Dale fellowship from the Wellcome Trust and Royal Society (213455/Z/18/Z) and a Lord Kelvin/Adam Smith (LKAS) Fellowship from the University of Glasgow. C.J.T. is supported by NHMRC in Australia (GNT1183496). ML.W. is supported via Australian Research Training Program Scholarship. J.G.W. is supported by a startup grant from the School of Life Sciences and College of Liberal Arts and Sciences at Arizona State University. E.S.MD is supported by CNPq (408964/2018-9) and CAPES grants. We thank Glasgow Polyomics

## Figures and legends

**Figure S1.**
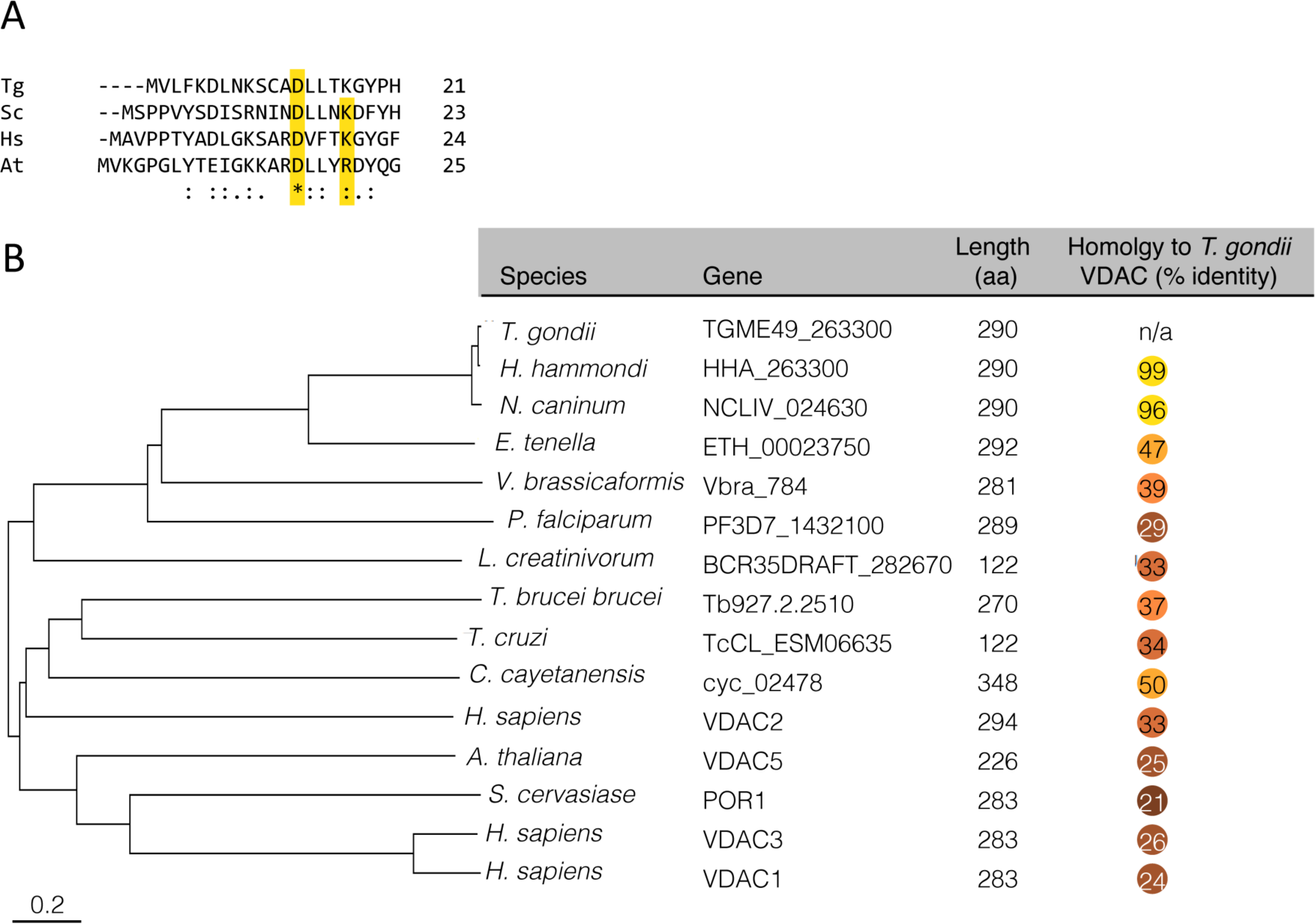
Phylogenetic analysis of VDAC from *T. gondii*. **A)** VDAC from *T. gondii* (Tg) contains glycine and lysine residues at positions 16 and 20 respectively which are conserved between human (Hs), yeast (Sc) and plants (At, D21 only). **B)** Molecular phylogenetic analysis using Neighbor-Joining method. Table showing the gene ID, length (in amino acids) and percentage identity to *T. gondii* of VDACs from selected species.

**Figure S2.**
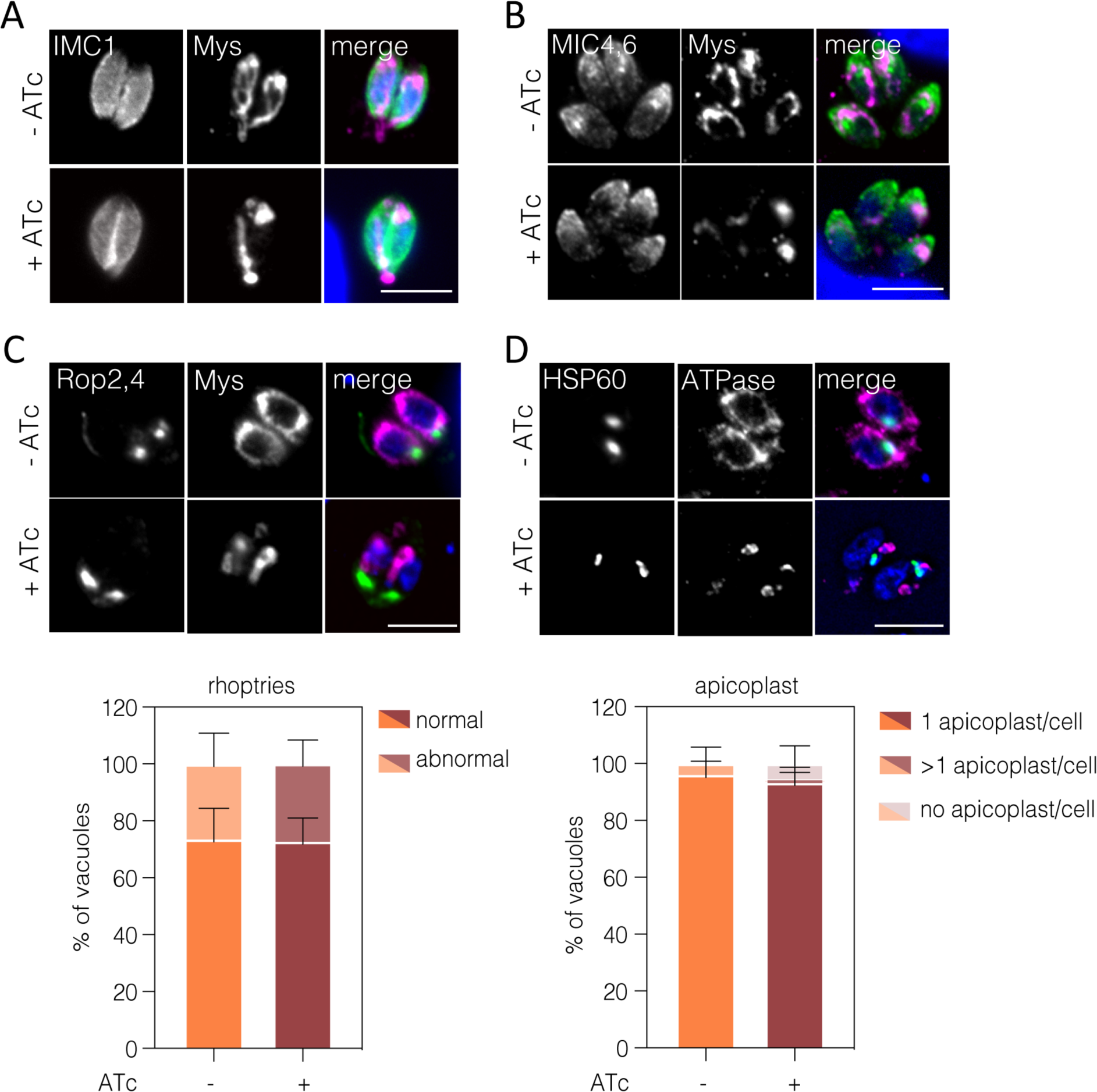
Depletion of VDAC does not affect other organelles besides mitochondrion and ER. iVDAC parasites ATc treated for 48 h were fixed and stained using markers for various organelles. No difference in parasite structure (from at least 260 parasites), as visualized by anti-IMC1 **(A)** or micronemes (from at least 174 parasites), visualized using anti-MIC4, 5 **(B)** could be seen upon VDAC depletion. Imaging and quantification of the rhoptries **(C)** using anti-ROP2, 4 and the number of apicoplasts **(D)** using anti-HSP60. No significant differences could be seen in either organelle. Scale bar 5 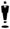m.

**Figure S3.**
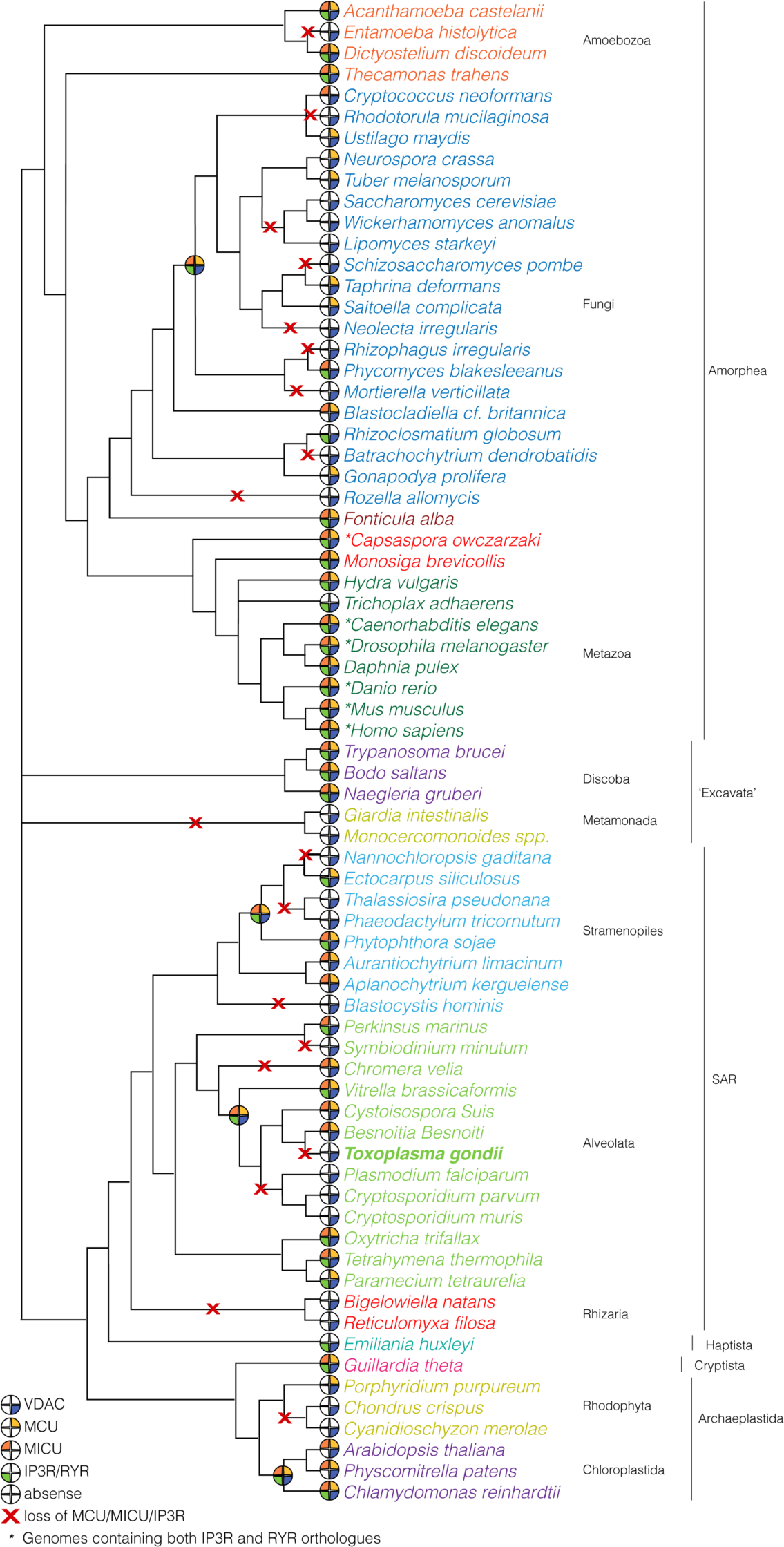
MCU, MICU, and IP3R have been lost 19 times in the evolution of eukaryotes. Orthologues of MCU, MICU, and IP3R were identified using BLAST and phylogenetic reconstructions. Organisms with identified orthologues are marked as indicated in the figure legend. Most eukaryotes contain homologues that could not be unambiguously identified as ITPR1 or RYR3. Asterisks indicate organisms containing both ITPR1 and RYR3 orthologues.

## Materials and Methods

### Parasite culture and genetic manipulation for line generation

*T. gondii* parasites were grown in human foreskin fibroblasts (HFFs) cells in supplemented Dulbecco’s modified Eagle’s medium (DMEM) supplemented with Penicillin/Streptomycin with 10 % of foetal bovine serum (FBS) (Gibco). Parasites were incubated at 37 °C with 5 % CO2 and 100 % humidity.

For transfections, electroporation was performed using 1 × 10^7^ freshly egressed parasites using a BioRaD electroporator following the manufacturer’s instructions. To localize tagged Myc-VDAC, parental TATiΔku80 (Sheiner et al., 2011(Jacot et al., 2014) parasites were transfected with a plasmid containing a pTUB8 promoter and VDAC cDNA coding sequence fused with an N-terminal Myc epitope tag, amplified using primers P1 and P2 (Table S1), named, pTUB8_Myc_VDAC.

To create the iVDAC parasite line, the promoter replacement strategy was used following a strategy previously described (Jacot et al., 2014; Sheiner et al., 2011). The ChopChop tool was used to identify the best gRNA containing the ATG of the gene (http://chopchop.cbu.uib.no/) and primers P3 and P4 were duplexed. gRNAs were cloned into a gRNA and Cas9 expression vector (Addgene #80636) using the BsaI restriction site. An PCR product containing the ATc repressible promoter followed by DHFR selectable cassette was amplified by PCR from pDT7S4myc using primers in P5 and P6 (**Table S1**) (Jacot et al., 2014; Sheiner et al., 2011). Parasites were co-transfected with 50 μg of the gRNA/CAS9 vector and the purified PCR products into the TATiΔku80 parental line. Cassette integration was selected for using 1 µM pyrimethamine for 3 weeks. After cloning into 96 well plates, clones were PCR screened using primers P7 and P8 (5’ integration) and P9 and P10 (3’ integration) (see **Table S1**) and sequenced.

To generate the tdTom/iVDAC line, pCTR2T-TGME49_215430-tomato (tandem tomato (2t) expression cassette) (van Dooren et al., 2016) transgene was introduced into the iVDAC line, followed by enrichment of the stably expressing fluorescence parasite population by cell sorting. Clones of this line were isolated by serial dilution into 96 well plates and fluorescent clones selected.

In all experiments, 0.5 µM anhydrous tetracycline (ATc) was added to iVDAC line to repress the regulatable promoter.

### Identification of TgVDAC and phylogenetic analyses

The bioinformatics tool tBLASTn was used to compare amino acid sequences against the translated nucleotide *Toxoplasma gondii* database to find a potential candidate. For the phylogenetic tree, the evolutionary history was inferred using the Neighbor-Joining method (Saitou and Nei, 1987). The optimal tree with the sum of branch length = 14.07947748 is shown. The tree is drawn to scale, with branch lengths in the same units as those of the evolutionary distances used to infer the phylogenetic tree. The evolutionary distances were computed using the Poisson correction method and are in the units of the number of amino acid substitutions per site. The analysis involved 15 amino acid sequences. All positions containing gaps and missing data were eliminated. There were a total of 226 positions in the final dataset. Evolutionary analyses were conducted in MEGA6 (Tamura et al., 2013).

### Immunofluorescence assay

Parasites were inoculated onto 24 well plates containing HFF cells on coverslips for the different time points, washed with phosphate buffered saline (PBS) and cells were fixed in 4% paraformaldehyde for 20 minutes at room temperature (RT). After three PBS washes, cell membranes were permeabilized and sample blocked by incubation with 0.2% Triton X-100/PBS (PBST) and 2% bovine serum albumin (BSA) for 20 min at RT. Cells were then incubated for 1 hour with the correspondent primary antibodies at RT in a wet chamber. Primary antibodies were diluted in 1% BSA in PBS-Triton (**Table S2**). Cells were washed three times with PBST and incubated with the correspondent secondary antibody (diluted in 1% BSA in PBST) (goat anti-mouse or anti-rabbit conjugated to AlexaFluor 594 or 488 1:1,000 (Invitrogen)) for 1 hour at RT in the dark. Following final washes, nuclei were stained by addition of 1 µg/mL 4’,6-diamidino-2-phenylindole dihydrochloride (DAPI; Sigma-Aldrich) containing FluoromountG (Southern Biotech). Images were acquired using a Delta Vision microscope as described (Ovciarikova et al., 2017) and were analysed using FIJI Image J 64 software (Schindelin et al., 2015).

### Quantitative RT-PCR (RT-qPCR)

For testing downregulation of VDAC gene upon ATc addition at different time points, RT-qPCR was performed as previously described (Biddau et al., 2018). Total RNA was isolated from freshly egressed treated parasites using TRIzolTM (Invitrogen, Thermo Fisher Scientific) following manufacturer’s instructions. To remove DNA contamination in the sample, RNA was treated with RNase-free DNase (Invitrogen, Thermo Fisher Scientific). 0.5 µg of total RNA was used to synthesize cDNA using SuperScript® VILO™ cDNA Synthesis Kit (Invitrogen, Thermo Fisher Scientific) following manufacturer’s instructions. qPCR was completed on a real-time PCR instrument (7500 Fast Real-Time PCR System, Applied Biosystems) with the Applied BiosystemsTM PowerUPTM SYBRTM Green Master Mix from Applied Biosystems using primers P11 and P12 (**Table S1**).

### Parasite growth assay

Freshly egressed tdTom/iVDAC parasites were filtered through a 3-μm polycarbonate filter, centrifuged, and resuspended in parasite culture medium without phenol red (Gibco BRL Life Technologies, Rockville, Md.). 500 parasites/well were used to infect confluent HFF cells in black optical bottom 96-well culture plates in triplicate. Parasites were treated (or not) with 0.5 µM ATc and fluorescence was read daily in a BMG Fluostar plate reader with constant gain, and data from the 3 wells were averaged. Blank wells, containing media and host cells with ATc were measured simultaneously on the same experimental plate. Parasite growth was evaluated by increase of fluorescence over time, normalized to day 0.

### Mitochondrial potential using JC-1

Freshly lysed parasites were 3.0 µm-filter purified and incubated with 1.5 µM of JC-1 (ThermoFisher Scientific) for 15 min at 37°C. After incubation, parasites were washed three times by centrifugation and re-suspended in DMEM without phenol red (Gibco BRL Life Technologies, Rockville, Md.). Treatment with 10 µM valinomycin (Val) for 10 min was included as a depolarising control. Unstained controls were used to define gates for analysis. 50,000 events per treatment, were collected on BD FACSCalibur^TM^ (Becton Dickinson, San Jose, US) flow cytometer and data were analysed using BD CellQuestPro for changes in the ratio of green to red, as an indicator of fluctuations in membrane potential.

### Protein Import Assay

Parasites were transfected with Hsp60L-mDHFR-cMyc (van Dooren et al., 2016) (kind gift from Giel van Dooren), 24 hours prior to the desired time point after growth in the presence or absence of 0.5 *µ*M ATc. Freshly egressed extracellular parasites were filtered, collected by centrifugation (1,500rpm, 10 min), washed twice with PBS and pellets were lysed in 800 *µ*L ice-cold lysis buffer NuPAGE LDS (Invitrogen) with 200 *µ*L methanol and 2% β-mercaptoethanol. Samples were boiled for 5 min at 95 °C. 10 μg of total protein, quantified using nanodrop, were separated on 12% SDS-PAGE gel and transferred (transfer buffer: 0.025 M TRIS, 0.192 M glycine, 10 % methanol) to a nitrocellulose membrane (0.45 μm, Protran™) for 60 minutes at 100 V. To control for protein transfer, the membrane was stained with Ponceau S red staining solution (Sigma-Aldrich) after transfer. The membrane was then blocked with 5% bovine serum albumin (BSA) in Tris-buffered saline (TBS; 20 mM Tris, 150 mM NaCl) and 0.05% Tween-20 (Sigma-Aldrich) and incubated with the primary antibodies (**Table S2**) overnight at 4°C in a wet chamber. After three washes with TBS; anti-mouse horseradish peroxidase (HRP)-conjugated secondary antibody (Promega) (1:10,000) was incubated at RT for 45 min with agitation. The signal was developed using Pierce^TM^ ECL Western Blotting Substrate (Thermo Scientific). FIJI was used for analysis of band intensity and the ratio between the mature and premature band intensity was quantified.

### Mitochondrial morphology analysis

Mitochondria were visualized by immunofluorescence as described above. 50 vacuoles of each treatment were counted and mitochondrial morphology was assessed following the criteria: six different mitochondrial morphology categories were assigned: (a) “open lasso”, (b) “lasso” (c) “connected”, (d) “sperm”, (e) “broken” and (f) “ball-like”. Parental and iVDAC parasites were grown in the presence or absence of ATc 0.5 µM for 24, 48 and 72h and mitochondria were visualized by IFA using an anti-TgMys antibody, as described above. Experiments were performed in triplicate.

### Super-resolution Fluorescent Microscopy

For super-resolution structural illumination microscopy (3D-SIM), stacks of vacuoles containing two or four parasites (with increments of 0.1µm in a total of 5 µm) were imaged in a Zeiss Elyra PS.1 super-resolution microscope (Jena, Germany) with a 63x/1.4 oil-immersion objective using ZEN black software (Zeiss, Germany). 5-phase SR-SIM images were reconstructed in the same software using Structural Illumination manual processing tool. 3D models were reconstructed in Imaris software (Oxford Instruments). The same software was used to calculate the volume and surface area of mitochondrial and ER signals. Co-localization tool in the same software was used to calculate the overlap proportion (Meanders coefficient calculation) of mitochondrion:ER signals.

### Transmission electron microscopy and morphometric analysis

LLC-MK_2_ cultures in 25 cm^2^ flasks were infected with tachyzoites of iVDAC mutant and incubated (or not) with 0.7 *µ*M ATc for 72 hours. After that, infected cells were fixed with 2.5 % glutaraldehyde in 0.1 M sodium cacodylate buffer (pH 7.4) and post-fixed for 45 min in the dark in 1% osmium tetroxide, 1.25% potassium ferrocyanide and 5 mM CaCl_2_, in 0.1 M sodium cacodylate buffer (pH 7.4). After post-fixation, infected cells were *en bloc* stained with uranyl acetate and lead aspartate then samples were dehydrated in acetone solutions of increasing concentrations (30-100%) and embedded in PolyBed 812 resin (Polyscience Inc., Warrington, PA, USA) using flat-embedding molds (EMS, Hatfield, PA). Ultrathin sections (70-80 nm) from three different blocks of -ATc and +ATc iVDAC were obtained in a Leica UC6 ultramicrotome and collected in 400 mesh copper grids (2 grids per block). Sections were observed in a JEOL 1200 EX and FEI Tecnai Spirit 120 transmission electron microscope and obtained images were analysed using Image J software.

### Gliding assay

Freshly egressed tachyzoites were 3.0 µm-filter purified, spun down and resuspended in Hank’s balanced salt solution supplemented with 100 mM HEPES (HBSS-H), 20mM EGTA at around 5 × 10^6^ parasites/mL. All the assays were done in HBSS not containing Mg or Ca, unless indicated. Parasites were layered onto poly-L-lysine-coated coverslips and allowed to adhere for 5 - 15 min at RT then incubated for 20 min at 37 °C, in the presence or absence of 2 mM ionomycin (Santa Cruz Biotechnology, 56092-82-1 Ionomycin-3592). After incubation, parasites were fixed as above followed visualization of gliding trails using the α-SAG1 (**Table S2**) with no permeabilisation. Images were acquired using a Delta Vision microscope as described (Ovciarikova et al., 2017). Image brightness was adjusted using FIJI software to allow for clear visualisation of the trails which were then traced with a wand tool and length of the trail was measured.

### Invasion assay

Parasite invasion was assessed using an invasion assay. 5 × 10^6^ artificially egressed parasites were centrifuged for 5 min at 500 x g then resuspended in 200uL volume and inoculated on confluent HFF cells on glass coverslips in a 24 well plate and incubated for 20 min at 37 °C before washing with PBS and fixing. Extracellular parasites were stained using anti-SAG1 antibody (**Table S2**) in non-permeablilized cells. Subsequently, cells were permeabilized and stained with rabbit anti-GAP45 (**Table S2**) to visualise all parasites. A minimum of 200 parasites were counted from 20 fields per treatment, for each experiment and the ratio of intracellular to extracellular parasites calculated.

### Metabolomics extraction

Stable isotope labelling of intracellular *T. gondii* tachyzoites was done as described below with some modifications from previous protocols described for *T. gondii* (MacRae et al., 2012; Oppenheim et al., 2014). 2 × 10^8^ intracellular parasites were washed with glucose-free DMEM containing glutamine, FBS and antibiotics. That media was immediately replaced with cold, glucose-free DMEM containing 4 mM ^13^c-U glucose (50:50). Parasites were incubated for 4 hours under standard culture conditions and harvested. Both host cell and parasite metabolism was quenched by placing the culture flasks on ice. Cells were washed twice and the parasites scraped, syringed and filtered to remove host cell debris into ice cold PBS. Parasites were pelleted at 4,000 rpm, for 25 min at 4°C. After washing the pellet was used to extract the sample in chloroform/methanol/water (1:3:1 v/v) buffer. After sonication for 2 min on ice, samples were incubated on ice under shaking for an hour with sonication every 10 min. Samples were then centrifuged at 13,000 g for 10 min at 4 °C. Supernatant was stored at 80°C prior to Liquid chromatography–mass spectrometry (LC-MS) analysis.

### Liquid chromatography-mass spectrometry analyses

Metabolomics analyses were performed by liquid chromatography-mass spectrometry using an Ultimate 3000 LC system (Dionex, UK) connected to a Q Exactive HF Hybrid Quadrupole-Orbitrap mass spectrometer, (Thermo Fisher Scientific). The system was controlled by the software Chromeleon (Dionex, UK) and Xcalibur (Thermo Scientific), acquiring both positive and negative ionisation mode. Chromatographic separation was performed with a ZIC-pHILIC chromatography column (150 mm 64.6 mm 65 mm; Sequant, Uemå, Sweden) using a two solvent system consisting of solvent A: 20 mM ammonium carbonate and solvent B: acetonitrile. The table shows chromatographic conditions:

| Solvent A (%) | Solvent B (%) | Time (min) |
| --- | --- | --- |
| 20 | 80 | 0 |
| 80 | 20 | 30 |
| 95 | 5 | 31 |
| 95 | 5 | 35 |
| 20 | 80 | 36 |
| 20 | 80 | 46 |

### Metabolomic data analysis

Vendor-specific raw data were initially centroided and converted into the open format mzXML for subsequent processing. PeakML files (Scheltema et al., 2011) were then generated by extracting the chromatographic peaks contained in the mzXML files using the detection algorithm from XCMS (Tautenhahn et al., 2008). The data processing pipeline mzMatch.R (Jankevics et al., 2012) was used to sort and combine all PeakML files corresponding to replicates and to exclude all non-reproducible data. Further steps of noise-filtering, gap-filling, and metabolite identification were performed on PeakML files utilising data obtained from metabolic standards run in parallel. For each metabolite of interest, the proportions of each isotopologue and its relative abundance in the sample were determined. The PeakML.Isotope.TargetedIsotopes function of mzMatch-ISO (Chokkathukalam et al., 2013) was used to scan the PeakML files for labelled metabolite quality and quantity.

### Statistical analysis

Data were analysed using GraphPad Prism software (version 5.00, La Jolla, US). Unless otherwise indicated, analyses were performed using either Student’s t-test or one-way ANOVA test with correction for multiple comparisons as indicated.

### Cytosolic [Ca2+] quantification

GCaMP6 and mCherry expressing plasmid (described in Stewart et al 2017) was linearised with the uprt homology region and transfected into tachyzoites and selected with FUDR. Stable parasite populations were then treated with and without ATc and GFP and mCherry fluorescence measured using FACS LSR IIW as described in Uboldi et al 2018 PLoS Biol. Briefly, A23187 and BIPPO was serially diluted at a 2x concentration. Basal tachyzoite fluorescence was measured and equivolume of 2x agonist was added rapidly before further data acquisition for 2 minutes. All analysis was performed in FloJo v10.

### Homology searching and phylogenetic reconstruction

MCU, MICU, IP3R, and RYR3 homologues were identified by using human sequences as BLAST queries into a subset of predicted proteomes from across the eukaryotic tree of life (**Table S3**) using the NCBI BLAST server, and the Joint Genome Institute’s Mycocosm and Phycocosm databases (Grigoriev et al., 2014). Potential homologues were validated by a reciprocal BLAST into the *Homo sapiens* predicted proteome. When the original *H. sapiens* sequence was recovered, we concluded that an orthologue was identified. Validated orthologues were then used to identify extremely divergent orthologues in some lineages (e.g., apicomplexans). Accession numbers of retrieved sequences can be found in **Table S3**. To ensure that no highly divergent sequences were overlooked, we used our validated orthologue set to generate an Hidden Markov Model via the HMMer server at EBI (Finn et al., 2011). We were unable to identify any additional candidate orthologues within our target set of species. We identified 19 instances where MCU, MICU, and IP3R were lost and mapped these losses onto a consensus tree of eukaryotes (**Figure S3**).

## Tables

**Table 1.**
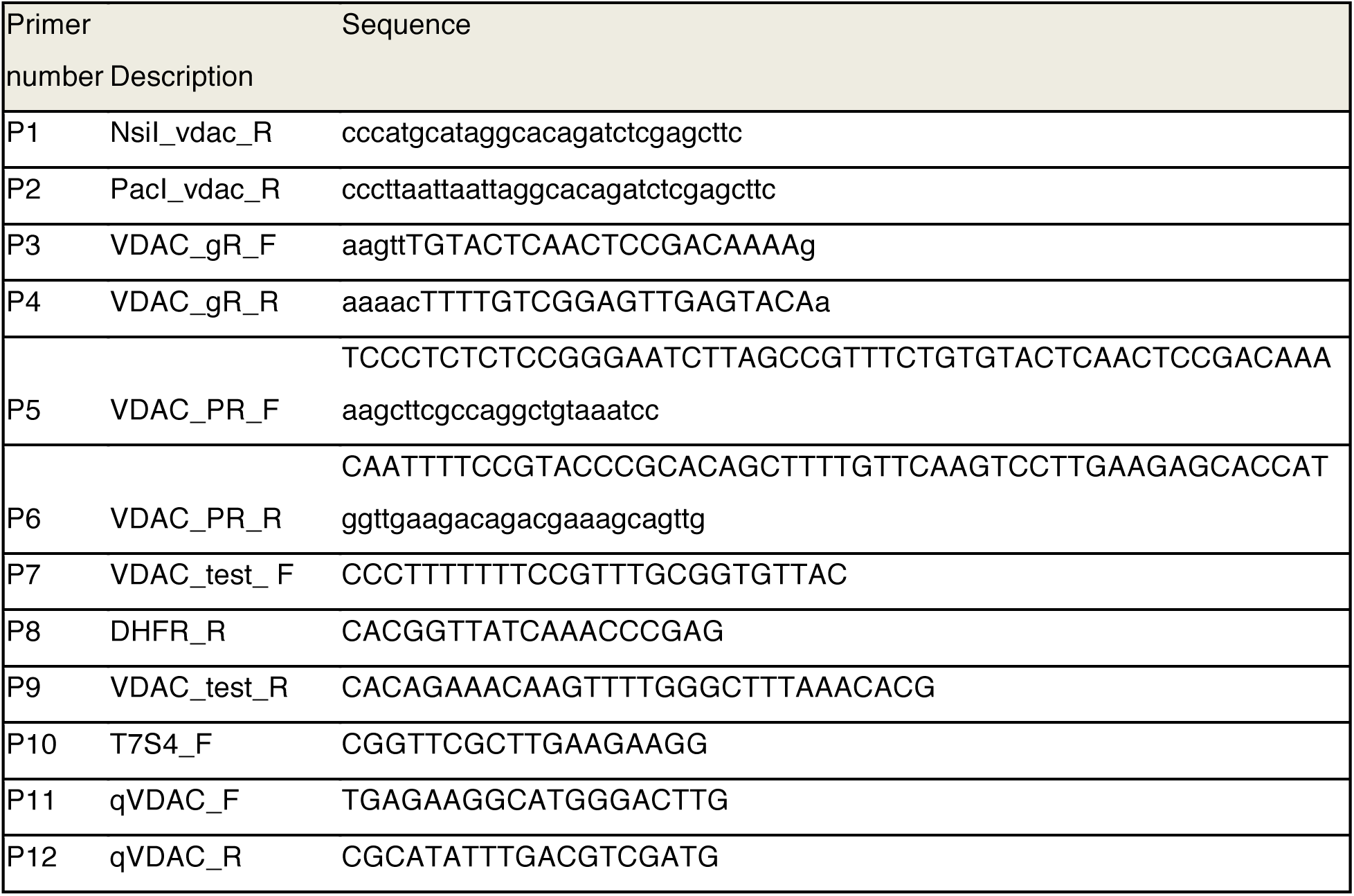
List of primers used in this study.

**Table 2.** List of antibodies used in this study.

| Antibody | Species | Dilution (IFA unless indicated) | Source | Reference |
| --- | --- | --- | --- | --- |
| CPN60 (HSP60) | Rabbit | 1:1000 | Boris Striepen (University of Pennsylvania) | (Agrawal et al., 2009) |
| IMC1 | Rabbit | 1:2000 | Gary Ward (University of Vermont) | (Wichroski et al., 2002) |
| Mys (TgME49_215430) | Rabbit | 1:1000 |  | (MacRae et al., 2012) |
| Myc | Mouse | 1:1000 | Cell Signaling |  |
| Myc | Rabbit | 1:1000 | ThermoFisher |  |
| GAP45 | Rabbit | 1:10000 | Dominique Soldati (University of Geneva) | (Plattner et al., 2008) |
| Sag1 | Mouse | 1:1000 | Sebastian Lourido (Whitehead Institute) | (Burg et al., 1988) |
| ROP2,4 | Mouse | 1:500 |  | (Sadak et al., 1988) |
| F1-ATPase | Mouse | 1:3000 | Peter Bradley (University of California, Los Angeles) | (Beck et al., 2014) |
| MIC4 | Rabbit | 1:5000 |  | (Reiss et al., 2001) |
| Actin | Mouse | 1:100 | Markus Meissner (Ludwig-Maximilians University of Munich) | (Angrisano et al., 2012) |

